# Associated organisms inhabiting the calcareous sponge *Clathrina lutea* in La Parguera Natural Reserve, Puerto Rico

**DOI:** 10.1101/596429

**Authors:** Jaaziel E. García-Hernández, Nicholas M. Hammerman, Juan J. Cruz-Motta, Nikolaos V. Schizas

## Abstract

Sponges provide an array of ecological services and benefits for Caribbean coral reefs. They function as habitats for a bewildering variety of species, however limited attention has been paid in the systematics and distribution of sponge-associated fauna in the class Calcarea or for that matter of sponges in the Caribbean. The goal of this study was to characterize infaunal assemblages from a calcareous sponge, *Clathrina lutea*, across multiple reefs from the La Parguera Natural Reserve, Puerto Rico. The associated fauna from 43 *C. lutea* specimens yielded a total of 2,249 associated infauna distributed in seven invertebrate phyla. Arthropoda was the most abundant phylum accounting for 62.5% of total abundance, followed by Annelida (21.0%) and Nematoda (5.5%). Limited patterns of temporal or spatial variability were surmised due to the opportunistic sampling effort afforded to this investigation from the cryptic nature of this species. A concordance between our data set and those for the class Demospongiae were observed, with the most abundant associated fauna being copepods and polychaetes. However, when compared to other Calcarea, the present study found considerably more associated fauna.

## INTRODUCTION

Marine sponges (Porifera, Grant, 1836) are perhaps the earliest metazoan phylum on the planet (Pisani et al. 2015), originating roughly 600 million years ago (Li et al. 1998; Yin et al. 2015). This evolutionary time has allowed sponges to develop complex biotic interactions with other marine organisms (Wulff 1985). Fossil evidence of associated phyla (Echinodermata, Bryozoa, Porifera, and Brachiopoda) in marine sponges from the early Ordovician Period has shed light into these ancient biotic interactions (Carrera 2000). Like their ancient counterparts, modern sponges provide habitat for an array of taxonomically diverse populations of micro-invertebrates and fish (Wulff 2006; Marliave et al. 2009; Gloeckner et al. 2014). As sessile organisms, sponges provide their symbionts a continuous flow of water and food in the form of phytoplankton, organic and inorganic detritus and in some instances even sponge tissue (Pawlik 1983; Corredor et al. 1987).

Sponges have been called ‘living hotels’ (Pearse 1932; 1950; Gerovasileiou et al. 2016), ‘living islands’ (Villamizar & Laughlin 1991) and ‘microcosms’ (Uriz et al. 1992); where different levels of interactions between host and colonizers are occurring simultaneously, often dependent upon the aquiferous system and morphology of the sponge (Koukouras et al. 1992; 1996). Long (1968) summarized these relationships into four groups: (1) inquilinism, or lodging, within or upon the sponges, (2) co-existence of two organisms on the same substratum because of simultaneous growth, (3) predation or grazing; and (4) mutualism.

Most of the studies regarding marine sponges and their associated fauna have focused on the class Demospongiae; perhaps because 86% of the extant Porifera are Demospongiae (van Soest et al., 2016). Among the most significant reports concerning demosponge-associated fauna were those of Santucci (1922), Pearse (1932, 1950), Fishelson (1966), Long (1968), Pansini (1970), Sube (1970), Labate & D’Addaboo (1974), Rützler (1976), Peattie & Hoare (1981), Koukouras et al. (1985), Wendt et al. (1985), Voultsiadou-Koukoura et al. (1987), Koukouras et al. (1992), Klitgaard (1995), Koukouras et al. (1996), Duffy (1996), Magnino et al. (1999), Ribeiro et al. (2003), Skilleter et al. (2005), Abdo (2007), Palpandi et al. (2007), Huang et al. (2008), Greene (2008), Schejter et al. (2012), Sivadas et al. (2014), and Schönberg et al. (2015). Of the mentioned studies regarding the associated organisms of demosponges, the most represented phyla are Arthropoda, Annelida, Mollusca, Nematoda, and Echinodermata and the most abundant classes are Crustacea, Polychaeta, and Ophiuroidea.

The remaining three sponge classes have received little attention worldwide, with six studies for Hexactinellida (Kunzmann 1996; Beaulieu 2001, a, b; Schuchert & Reiswig 2006; Fiore & Jutte 2010; Kersken et al. 2014), three for Calcarea (Frith 1976; Padua et al. 2013; Ribeiro et al. 2016), and none for the class Homoscleromorpha. This lack of knowledge prohibits an understanding of how associated assemblages could differ among sponge classes. Since sponges’ act as bioengineers, in effect creating habitat for potentially thousands of organisms, studying the lesser known Porifera groups can shed light on the functional roles these sponges provide, especially since sponges are becoming the dominant benthic group in Caribbean reefs (Loh et al. 2015). Even though research in the Caribbean region has yielded one of the most interesting associations between snapping shrimps and demosponges (eusociality; Duffy, 1996), relatively few investigations have quantified total sponge-associated fauna (e.g. Pearse 1932; Westinga & Hoetjes 1981; Villamizar & Laughlin 1991). Only two previous studies (Frith 1976; Padua et al. 2013) have looked at the associated assemblages of calcareous sponges, both focusing on sponges belonging to the subclass Calcaronea. Although Ribeiro et al. (2016) did not explore the associated organisms of the calcareous sponge *Clathrina aurea*, they did provide unique and important ecological data regarding the influence of intra- and interspecific interactions in the abundance, growth, and lifespan of the calcinean *C. aurea*. To our knowledge, this study is one of the few to explore the associated fauna of sponges belonging to the class Calcarea (subclass Calcinea) and will set the foundation for future studies focusing on this group.

The objectives of this study were to (1) describe the composition and structure of the invertebrate assemblages associated with the calcareous sponge *Clathrina lutea* in La Parguera (southwest Puerto Rico), (2) and describe patters of spatial and temporal variation of those infaunal assemblages. The studied calcareous sponge is a new species, *C. lutea* (Fig. 1), which was recently described by Azevedo et al. (2017). The genus *Clathrina* (Gray, 1867) has an asconoid aquiferous system, one of the simplest organization among sponges, which means that all the cavities are lined by choanocytes (Klautau & Valentine 2003). In Puerto Rico, *C. lutea* inhabits sciophilous habitats, usually growing on cavern walls and ceilings, overhangs, small crevices, and sometimes exposed on reefs. *C. lutea* can also be found growing at the base of gorgonians (*Erythropodium caribaeorum* and *Plexaura flexuosa*), zoantharians (*Palythoa caribaeorum*), and underneath scleractinian coral colonies (*Montastraea cavernosa, Orbicella annularis*, and *O. faveolata*) (García-Hernández, pers. obs.) (Fig. 1).

**Figure 1.**
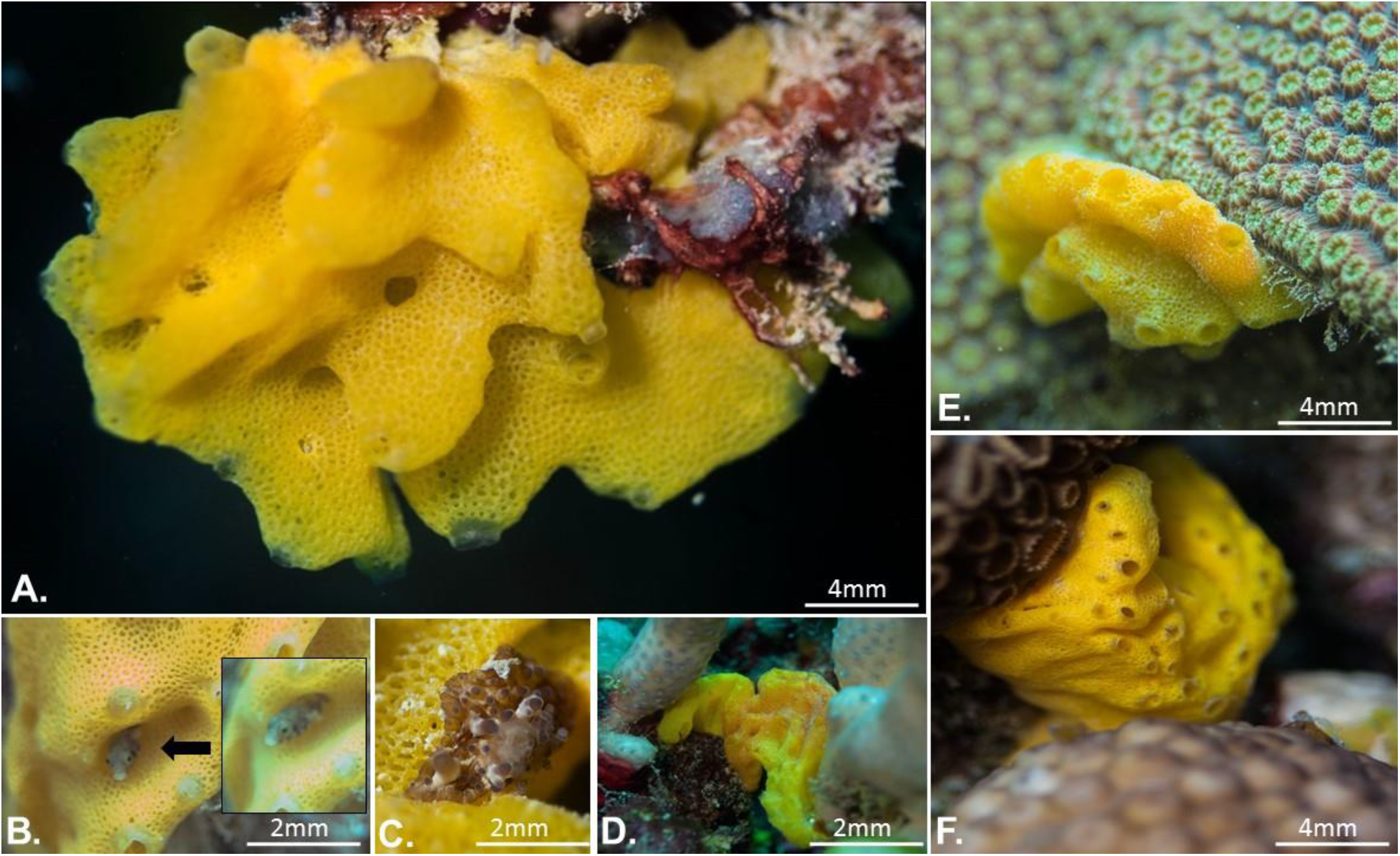
The Calcareous sponge *Clathrina lutea* A. *In situ* photograph of *C. lutea* growing in a cavern overhang, B. Juvenile fish taking refuge in the folds of the sponge, C. Sea-anemone, *Bunodeopsis* sp. attached to the sponge, D. *C. lutea* growing at the base of the octocoral, *Briareum asbestinum*, E. *C. lutea* growing underneath the scleractinian coral, *Orbicella faveolata*, and F. *C. lutea* growing underneath the white encrusting zoanthid, *Palythoa caribaeorum.*

## MATERIAL AND METHODS

### Sampling and Laboratory Dissections

*Clathrina lutea* is a cryptic species, whose patterns of spatial distribution are vaguely known, with reports in Brazil (Azevedo et al. 2017), Curaçao (Cóndor-Luján et al. 2018), and Puerto Rico (JEGH unpubl data). Consequently, a systematic sampling effort to ensure equally balanced replicates per site and time was not possible. Despite these challenges, a total of 43 specimens were collected from eight coral patch reefs off the southwest coast of La Parguera, Puerto Rico (Fig. 3) from April 2014 to April 2015 using SCUBA. Field collections consisted of wrapping a plastic bag around the sponge to reduce fast moving associated fauna escapement (mostly crustaceans) and then the sponge was placed into a large collection vial. In the laboratory, the sea water was discarded, and the remaining sample was labeled, fixed and preserved in 95% ethanol in a falcon tube. All sponges were stored in a −20 °C freezer for further analysis. Before dissection, wet weight was calculated per sponge, followed by examination of the ethanol in which the host was preserved. Sponge specimens were then carefully inspected and fragmented under a stereomicroscope to remove associated fauna from pores and canals as in Ribeiro et al. (2003). The associated fauna from both the ethanol and sponge were separated and identified to the lowest taxonomic level possible. Associated fauna were then preserved in new ethanol and vouchers deposited in the Caribbean Laboratory of Marine Genomics in the Department of Marine Sciences at the University of Puerto Rico at Mayagüez. Taxonomic resolution was performed down to the ordinal rank, with only few exceptions to the family level. This investigation describes the associated fauna within *C. lutea* at a coarse taxonomic resolution; allowing for overall comparisons between this study and others like it.

**Figure 2.**
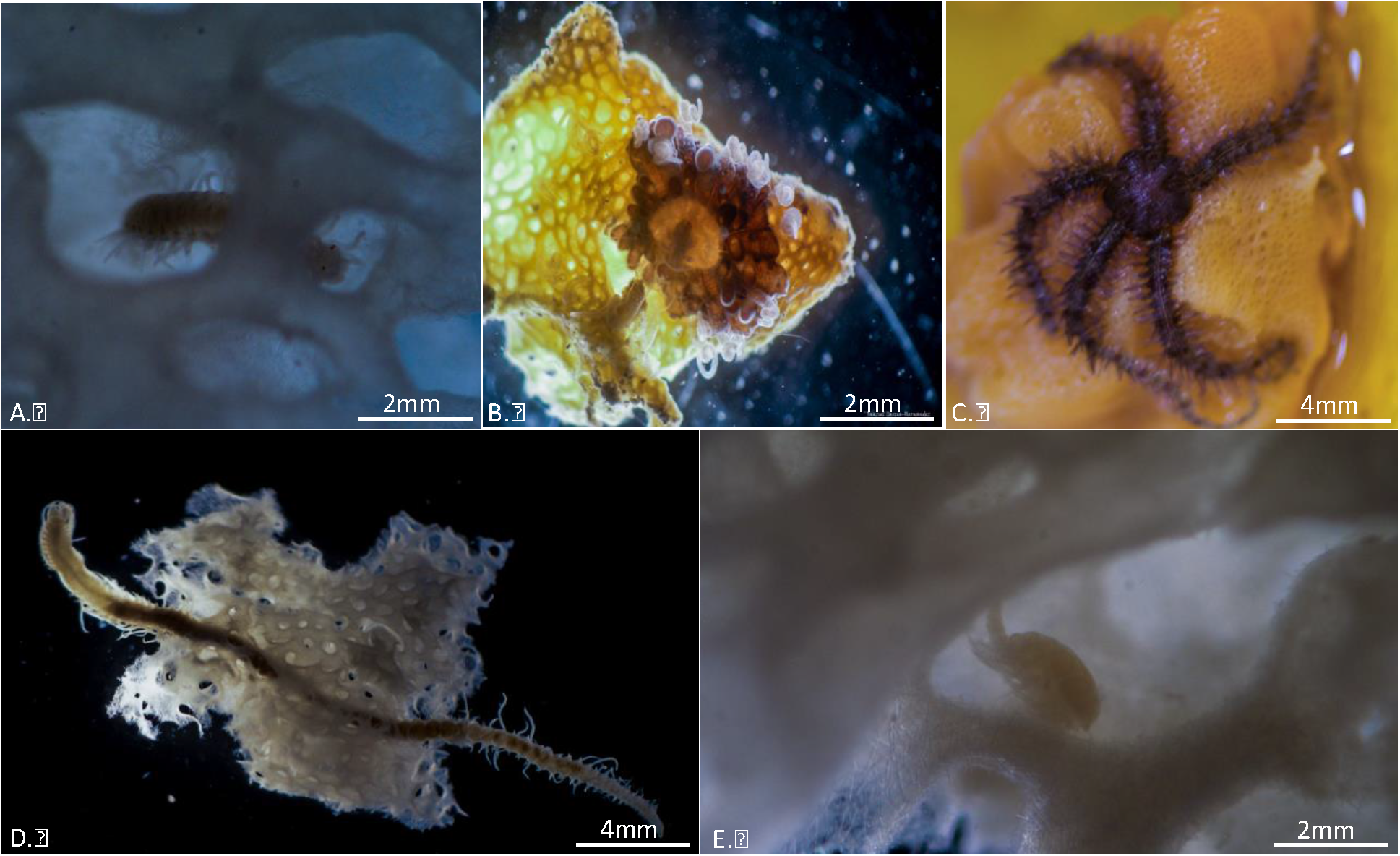
Associated infauna of the calcareous sponge, *Clathrina lutea*, A. Polychaete within the sponge, B. Sea Anemone, *Bunodeopsis* sp., with tentacles partially extended, C. Brittle star attached to the exposed internal canal, D. Polychaete found through a cross-section of the sponge and E. Copepod found within the interior cavity of *C. lutea*.

**Figure 3.**
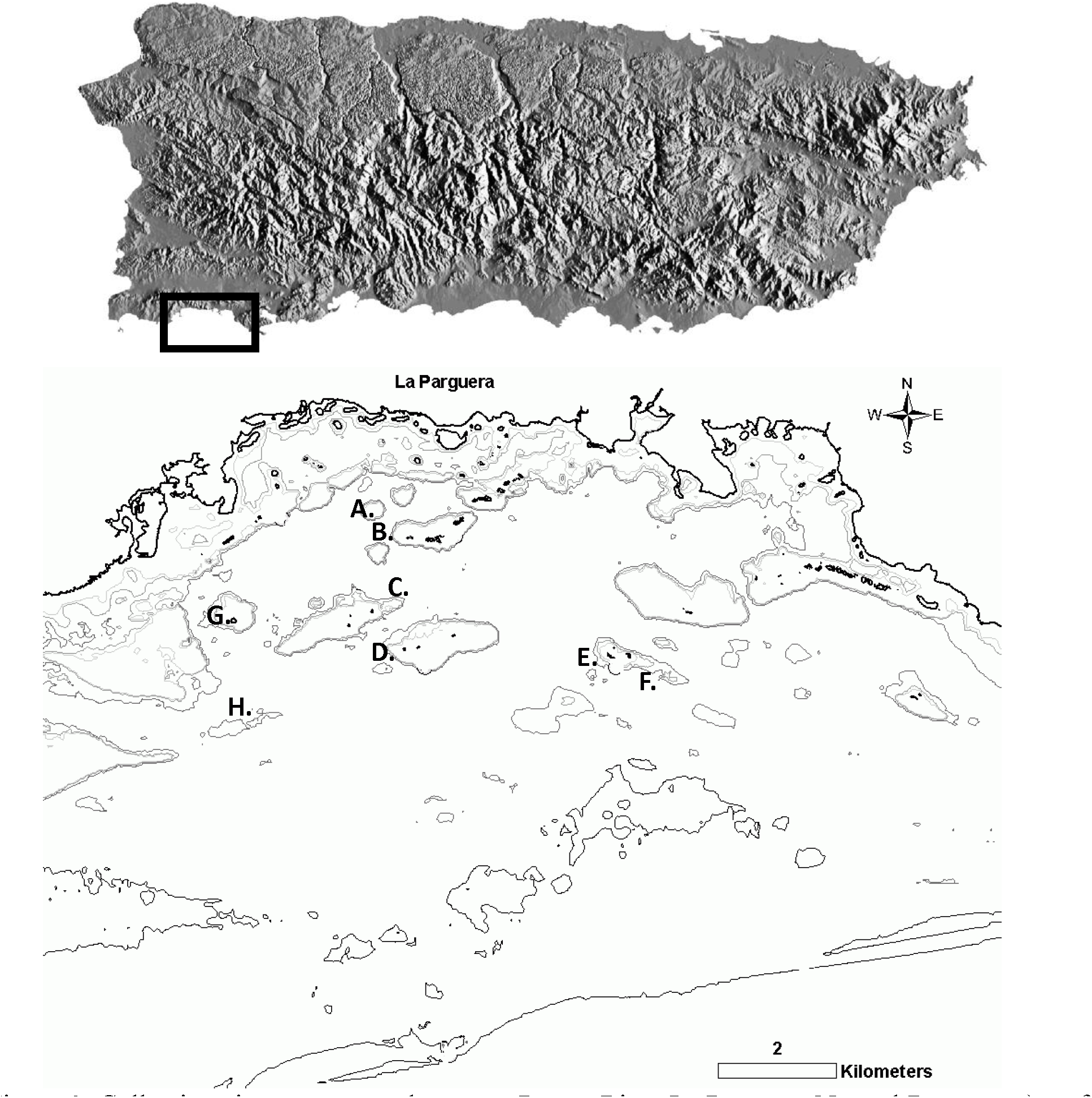
Collection sites across southwestern Puerto Rico, La Parguera Natural Reserve. a) reef Mario (17°95’N, −67°05’W), b) reef Enrique (17°57’N, −67°02’W), c) reef Laurel (17°56’N, −67°03’W), d) reef Media Luna (17°95’N, −67°05’W), e) reef Turrumote (17°93’N, −67°02’W), f) reef Pinnacles (17°90’N, −67°00’W), g) reef San Cristobal (17°93’N, −67°11’W), h) reef Margarita (17°95’N, −67°11’W).

### Data Analysis

Extraction and sorting of specimens resulted in data matrices for abundance of different invertebrate taxa per sponge that were used to construct similarity matrices among samples using the Bray-Curtis index. Before this, data was squared-root transformed to down-weight the dominance of highly abundant taxa in the calculation of similarities, relative to the less-common taxa. Also, data was standardized by total counts per sponge because size and volume of each sampling unit (i.e., sponge) was not the same. Decisions about transformations were done after visual inspections of Shade Plots (Clarke et al. 2014). Dissimilarity matrices were used to perform non-metric multivariate ordinations (nMDS) to illustrate patterns of spatial distribution of invertebrate assemblages associated with *C. lutea*. Given the nature of the sampling design used in this study, more detailed spatial-temporal analyses were done only for those sites that were sampled at least during two different seasons (i.e. the reefs Turrumote, San Cristobal and Pinnacles). In this case, Principal Coordinate Ordinations (PCO) were done on centroids per site and sampling time, which allowed interpretations of magnitude and direction of change among sites and across seasons. All analyses were done using the software PRIMER V7 (Clarke & Gorely 2015). Finally, correlations between sponge volume, number of individuals, and number of taxa of the associated infauna were calculated in relation to individual sponges.

## RESULTS

### Associated Infauna

The associated fauna from 43 specimens of the calcareous sponge *C. lutea* from eight coral patch reefs in southwestern Puerto Rico yielded a total of 2,249 associated organisms specimens in seven invertebrate phyla (Figs. 3 and 4, Table 1). Arthropoda was the most abundant Phylum accounting for 62.5%, followed by Annelida (20.6%), and Nematoda (5.47%).

**Table 1.** Total Counts of Associated fauna living within the sponge, *Clathrina lutea*, Art=Arthropoda, Cni=Cnidaria, Ann=Annelida, Ech=Echinodermata, Nem=Nematoda and Mol=Mollusca.

| Site |  | Mario | Enrique | Laurel | Media Luna | Turrumote | Pinnacles | Margaritas | San Cristobal | Total |
| --- | --- | --- | --- | --- | --- | --- | --- | --- | --- | --- |
| Phylum | Taxa |  |  |  |  |  |  |  |  |  |
| Art (7) | Isopoda | 14 | 2 | 10 | 23 | 27 | 20 | 1 | 15 | 112 |
|  | Tanaidacea | 0 | 1 | 0 | 0 | 2 | 4 | 0 | 0 | 7 |
|  | Amphipoda | 24 | 1 | 16 | 15 | 41 | 39 | 4 | 12 | 152 |
|  | Decapoda | 0 | 1 | 0 | 0 | 1 | 0 | 0 | 1 | 3 |
|  | Copepoda | 125 | 6 | 300 | 81 | 226 | 220 | 30 | 109 | 1097 |
|  | Ostracoda | 3 | 1 | 3 | 1 | 7 | 10 | 0 | 1 | 26 |
|  | Acari | 1 | 0 | 1 | 0 | 4 | 0 | 1 | 1 | 8 |
| Cni (1) | Leptomedusae | 1 | 10 | 3 | 4 | 4 | 4 | 0 | 2 | 28 |
| Bry (1) | Crisiidae | 1 | 0 | 11 | 8 | 9 | 14 | 0 | 12 | 55 |
| Ann (1) | Polychaetes | 44 | 41 | 30 | 46 | 139 | 64 | 14 | 86 | 464 |
| Ech (1) | Ophiuroidea | 4 | 0 | 6 | 5 | 8 | 2 | 0 | 23 | 48 |
| Nem (1) | Nematodes | 9 | 1 | 38 | 10 | 24 | 24 | 0 | 17 | 123 |
| Mol (2) | Gastropoda | 3 | 0 | 1 | 1 | 6 | 2 | 0 | 3 | 16 |
|  | Bivalvia | 0 | 3 | 0 | 1 | 6 | 6 | 0 | 3 | 19 |
| Other | Other | 5 | 6 | 19 | 13 | 19 | 13 | 1 | 15 | 91 |
|  |  |  |  |  |  |  |  |  |  | 2249 |

**Figure 4.**
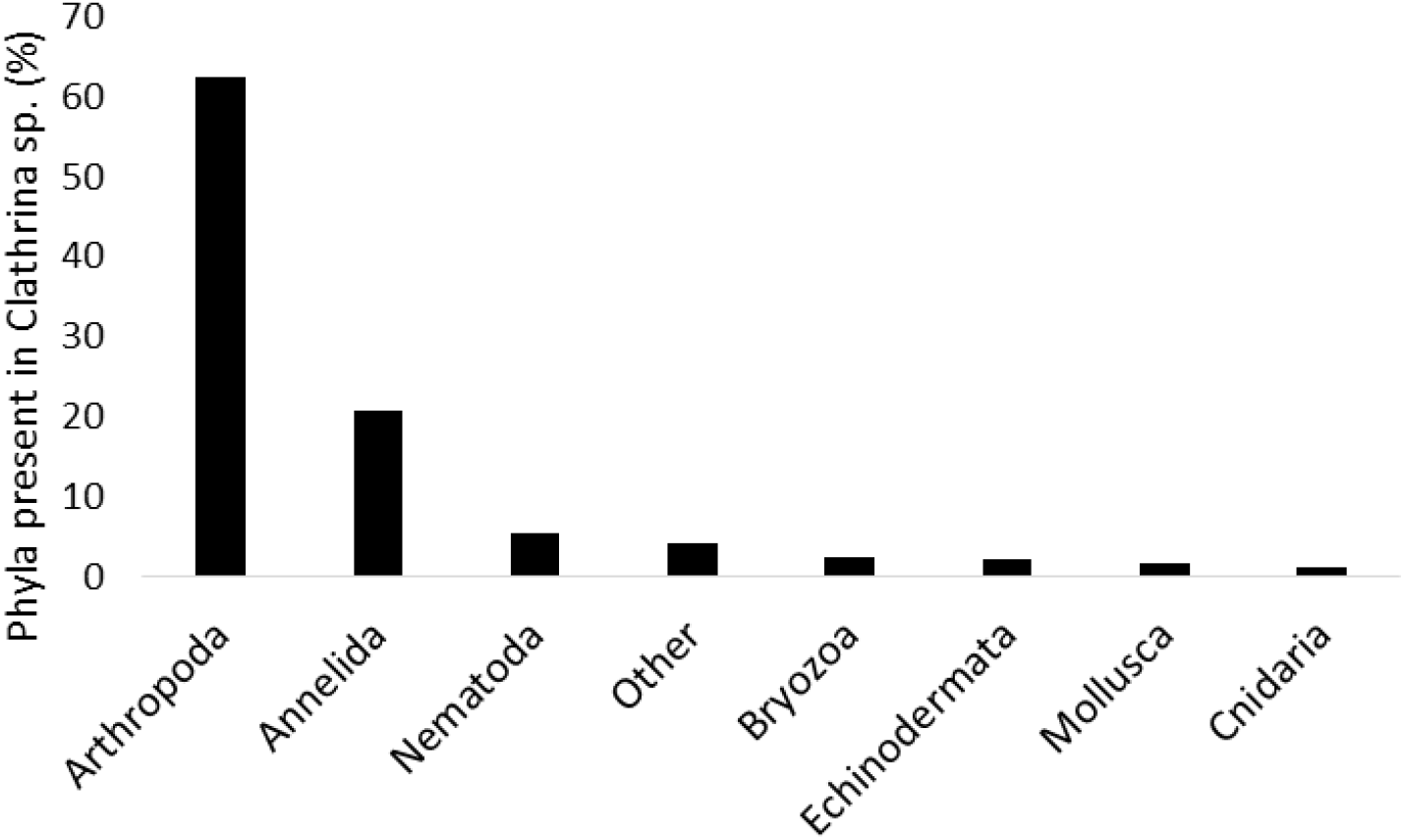
Invertebrate phyla of associated fauna found within the calcareous sponge *Clathrina lutea*.

Within the Arthropoda, the crustacean Copepoda yielded 48.8% of all associated individuals within *C. lutea.* The next most abundant groups within Arthropoda were the Amphipoda and Isopoda. The groups Tanaidacea, Decapoda, Ostracoda and Acari (halacarid mites) were also present (Table 1). The next largest group was the Phylum Annelida, all worms within the class Polychaeta and accounting for roughly 21.0% of total species abundance. The remaining phyla accounted for roughly 16.5% of taxa abundance, including Nematoda (5.5%), Bryozoa in the class Stenolaemata (2.4%), Echinodermata specifically the class Ophiuroidea (2.1%), Mollusca (1.6%) and the class cnidarian class Hydrozoa (1.2%). About 4% of total abundance were classified as unknowns (Fig. 4, Table 1).

### Reef site comparisons of infauna

Among all eight sites, the copepods, were the most abundant organisms found in *Clathrina* sp. (N=1,097, Table 1). Sponges dissected from Laurel reef contained the highest number of associated copepods (N=300), while Enrique reef contained the least (N=6). Decapods (N=3) were found in three different sponges from reefs Enrique, Turrumote, and San Cristobal. Tanaids (N=7) were found in reefs Enrique, Turrumote, and Pinnacles. Halacarid mites (N=8) were recovered from sponges collected from reefs Enrique, Laurel, Turrumote, Margaritas, and San Cristobal. Hydrozoans belonging to the order Leptomedusae and Nematodes were found in all reefs, except in sponges from Margaritas reefs. Bryozoans belonging to the family Crisiidae, as well as gastropods and ophiuroids were found in sponges from all reefs, except at Enrique and Margaritas. Polychaetes were found in sponges collected from all eight reefs. Microscopic bivalves were found in all reefs except at Mario reef and Margaritas reef. (Table 1).

Among all eight sampled reefs, twelve sponges from Turrumote reef contained the highest number of associated taxa (504 specimens belonging to 14 taxa), while two sponges from Margaritas reef contained the lowest number of associated taxa (53 specimens belonging to 7 taxa). Sponges collected from Laurel reef contained a relatively high number of associated taxa (19 specimens belonging to 11 taxa), even though only two specimens were found in this site. Interestingly, sponges collected from Laurel reef had the lowest average volume 1.41 cm^3^ (± 1.49, SD), yet their average density of individuals was the second highest of all sites with 46.7 ind.cm^-3^ (± 41.01, SD), while sponges dissected from Margaritas reef had the third highest average volume (2.95 ± 2.58 cm^3^), yet contained the lowest density of associated individuals with 13.3 ind.cm^-3^ (± 11.1, SD) (Table 2).

**Table 2.** Summary of ecological data collected at each site.

| Site | Number of<br>Taxa (S) | Number of<br>Specimens (N) | Average Volume<br>+/- SD (cm<sup>3</sup>) | Average Density<br>+/- SD (ind.cm<sup>-3</sup>) |
| --- | --- | --- | --- | --- |
| **San Cristobal**<br>**(n=9)** | 13 | 285 | 2.49 ± 1.87 | 16.8 ± 13.7 |
| **Margaritas**<br>**(n=2)** | 7 | 53 | 2.95 ± 2.57 | 13.3 ± 11.1 |
| **Media Luna**<br>**(n=3)** | 11 | 195 | 3.80 ± 0.698 | 17.0 ± 0.902 |
| **Mario (n=4)** | 11 | 229 | 3.35 ± 0.460 | 17.2 ± 2.041 |
| **Pinnacles**<br>**(n=9)** | 12 | 409 | 1.41 ± 1.49 | 49.1 ± 38.1 |
| **Turumote**<br>**(n=12)** | 14 | 504 | 2.50 ± 1.41 | 16.6 ± 15.3 |
| **Laurel (n=2)** | 11 | 419 | 3.18 ± 2.97 | 46.7 ± 41.01 |
| **Enrique**<br>**(n=2)** | 10 | 67 | 1.24 ± 1.58 | 33.9 ± 10.9 |

Overall, the average total volume of the sponges collected was 2.43 cm^3^ (±1.62). With respect to sponge volume, regression analysis indicated that, despite a positive slope, the average total volume of the sponges dissected did not significantly correlate with the number of individuals (N) (R^2^ = 0.215) and the number of taxa (S) (R^2^ = 0.230) (Fig. 5).

**Figure 5.**
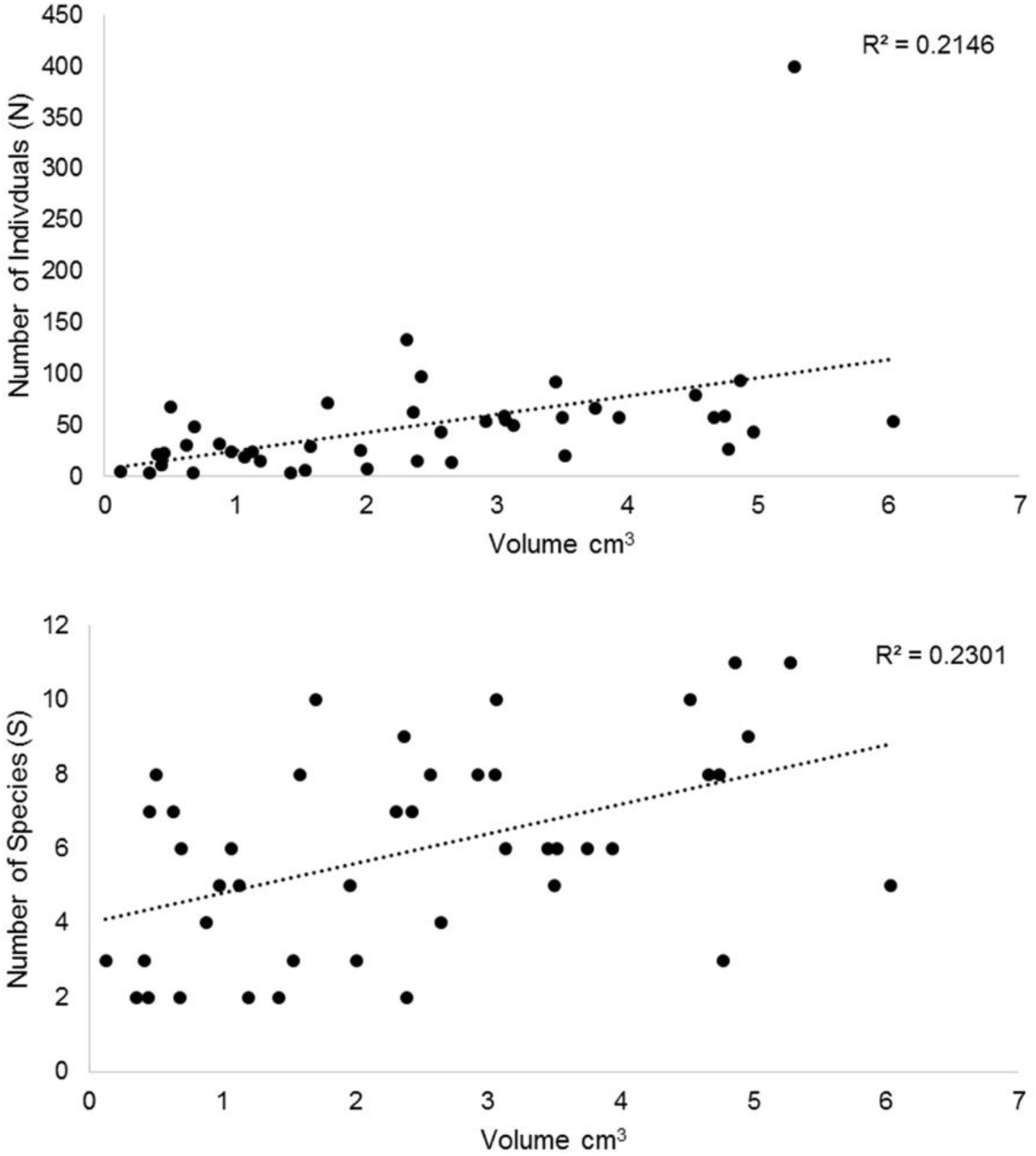
Linear regression between sponge volume and (A) number of individuals and (B) number of species of the infauna associated with *Clathrina lutea*.

### Temporal and Spatial Variation

Multivariate ordinations for assemblages of invertebrates associated with *C. lutea* showed no clear patterns of differences among reefs (Supplementary Fig. 1). This lack of patterns might be the result of important temporal variation as well as variation among sponges within a reef. To elucidate these patterns, multivariate ordinations (PCO) were done on centroids only for those sites that were sampled, at least, during two different seasons. These metric ordinations showed important temporal differences for San Cristobal reef, especially during the spring and fall of 2015, when polychaetes and ophiuroids increased in relation to other seasons in San Cristobal reef (Fig. 6).

**Figure 6.**
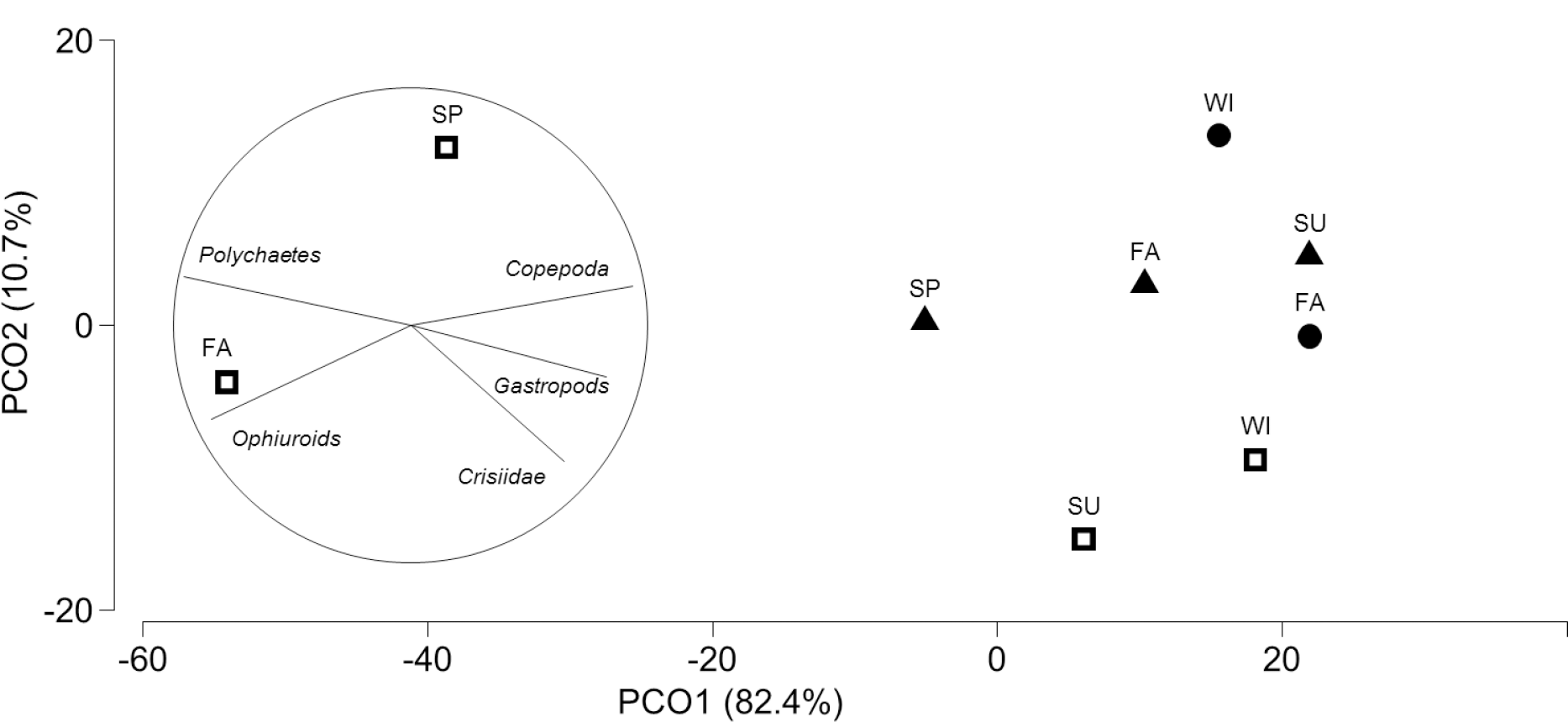
Principal Coordinate Ordinations (PCO) of centroids (site * season) of Bray-Curtis similarities calculated from data of invertebrate assemblages associated with *Clathrina lutea* in three different sites (▴ = Turrumote, ◻ = San Cristobal, • = Pinnacles) during four different seasons (SP = Spring, SU = Summer, FA = Fall, WI = Winter).

## DISCUSSION

After examination of 43 specimens of the small cryptic sponge, *C. lutea*, a myriad of associated organisms was classified into coarse taxonomic groups (Figs. 2 and 3), belonging to the Phyla Arthropoda, Annelida, Mollusca, Nematoda, and Echinodermata with the most abundant classes represented by Crustacea and Polychaeta (Table 1). The taxonomic resolution, to the ordinal rank yielded similar patterns and associated taxa between *C. lutea* and other sponges belonging to the class Demospongiae, with crustaceans being the most abundant (Pease 1950; Pansini 1970; Rützler 1976; Koukouras et al. 1985; Voultsiadou-Koukoura et al. 1987; Villamizar and Laughlin 1991; Koukouras et al. 1992; Ribeiro et al. 2003; Skilleter et al. 2005; Abdo 2007; Palpandi et al. 2007; Huang et al. 2008; Greene 2008; Schejter et al. 2012; Kersken et al. 2014; and Gerovasileiou et al. 2016). However, several other studies reported in Appendix 1 of Schejter et al. (2012) found Polychaetes to be the most abundant associated organisms.

The only other publication focusing solely on associated fauna in calcareous sponges discovered three additional phyla, Porifera, Platyhelminthes and Ascidians from *Paraleucilla magna* Klautau, Monteiro & Borojevic 2004, in Brazil (Padua et al. 2013). Surprisingly, of the 54 sponges examined, only 349 associated organisms were found in total within *P. magna*, on average 6.5 ind. specimen^-1^. As compared to this study, seven phyla were discovered from 43 specimens of *C. lutea* from southwestern Puerto Rico, with a total of 2,249 associated organisms, or 52.3 ind. specimen^-1^. Both species of sponges are the most conspicuous Calcarea in their respective geographical regions and can be found in similar photophilous and sciophilous environments (Klautau et at. 2004). The high number of associated fauna in the present study may be attributed to the differences in morphology between the two-sponge species as well as their prevalence through seasons. *P. magna*, belonging to the subclass Calcaronea, demonstrates strong seasonality throughout the year, being most abundant in summer and disappearing during autumn, only to reappear again in winter (Klautau et al. 2004). *P. magna* is also comprised of a leuconoid aquiferous system with large atrial cavities and numerous canals (Padua et al. 2013). *C. lutea* can be found readily year-round within the La Parguera Marine Reserve, belongs to the subclass Calcinea and is composed of an asconoid aquiferous system, with a cormus of anastomosed tubes consisting of a series of interwoven canals lined by choanocytes (Klautau and Valentine 2003). In addition, Frith (1976) surprisingly observed no associated fauna in three other species of Calcarea, belonging to the subclass Calcaronea from England. Thus, of the five species of Calcarea examined (Frith 1979; Padua et al. 2013; this study), *C. lutea* had the most associated individuals, which might be due to its prevalence through seasons as well as the difference in sponge’s internal morphology, allowing more space and crevices, thus more niches for the associated fauna. Interestingly, specimen dissections of *C. lutea* collected from Brazil did not yield associated organisms (M. Klautau pers. comm.). Although not tested in this study, this may be due to a difference in composition of allelochemicals between geographically distant populations of *C. lutea*. It has been previously shown that some of these chemicals can deter or maintain certain fauna within a species of sponge (i.e., Betancourt-Lozano et al. 1998).

Regression analysis yielded a positive but non-significant correlation between sponge volume and the absolute abundance of associated individuals and taxa (Table 2 and Figure 5). Past studies have shown similar results with respect to volume and number of associated individuals (Pansini 1970; Sube 1970; Koukouras et al. 1985; Fiore & Jutte 2010). Some specimens of *C. lutea* were particularly small but contained a high number of copepods, where other larger sponges often only had several organisms such as polychaetes and ophiuroids. In this case, the lack of a significant correlation between volume and total number of inhabitants, might be due to predation and competition. Previous literature has shown ophiuroids feeding on larvae of the Caribbean sponge *Callyspongia vaginalis* (Henkel & Pawlik 2014). Similarly, polychaetes have been documented as feeding upon sponge tissue and other smaller crustaceans (Pawlik 1983). Since many individuals of *C. lutea* had both brittle stars and polychaetes, they could limit the number of copepods, isopods, mites, and amphipods, if they were preying on them. Though within this study, gut content of associated fauna was not dissected.

In both *C. lutea* and in *P. magna* (Padua et al. 2013), Crustacea was the most abundantly represented group of associated organisms, 63% and 54%, respectively. Padua et al. (2013) found that amphipods within the family Stenothoidae, were the most abundant crustacean in the calcareous sponge *P. magna*. In comparison, specimens of *C. lutea* were dominated by copepods, with a total of 78% fauna abundance. Rützler (1976) also reports crustaceans (67.8%) as the most abundant group within six different species of sponges from the Mediterranean.

Interestingly, only within the sponge *Iricnia variabilis* where copepods the most abundant group, the remaining five were dominated by polychaetes, caridea, and gammaridea (Rützler, 1976). In the Aegean Sea, Koukouras et al. (1992) found that Crustacea was the most abundant taxon in four demosponges: *Agelas oroides, Petrosia ficiformis, I. variabilis*, and *Aplysina aerophoba*. In Brazil, Ribeiro et al. (2003) also reports crustaceans as the most abundant group (83%) within the sponge *Mycale microsigmatosa*. In the Mediterranean, Pansini (1970) found that copepods were the most abundant group within the demosponges *Spongia officinalis, Sarcotragus fasciculatus*, and *P. ficiformis*, 40.7%, 36.7%, and 36.1% respectively. Similarly, Westinga & Hoetjes (1981) report the presence of copepods in large quantities from 35 individuals of the demosponge *Spheciospongia vesparia*, however, these were not considered in their study due to difficulty in identification. Westinga and Hoetjes (1981) do note that most of their copepods were cyclopoid, of the genus *Asterocheres*, just like in this study. In deeper environments, Kersken et al. (2014) also found copepods to be the most abundant organism associated with the hexactinellid sponge *Rossella antarctica*.

When observing changes in associated assemblages across temporal or spatial variation, few clear patterns emerged, possibly due to the opportunistic sampling design which did not allow for equal collections across reefs through time (Supplementary Fig. 1). Of the reefs which were sampled evenly across seasons (i.e. Turrumote, San Cristobal and Pinnacles), only San Cristobal reef showed a transition from harboring predominately ophiuroids and polychaetes in the fall and spring to copepods and gastropods in the summer and winter (Fig. 6). It is possible that these temporal patterns of variation might be cyclical due to the predator/prey dynamics described above. Nevertheless, further studies on the trophic dynamics of assemblages associated with *C. lutea* will have to be conducted to unequivocally conclude this.

It would be difficult to partition associated fauna as either being facultative (i.e. actively select *C. lutea*) or ephemeral (i.e. transitory associates). However, the fact that multiple life stages of polychaetes were found in sponges across all eight reefs, suggests some of the associated fauna could be facultative. Especially for polychaetes since this was the second most abundant group within *C. lutea*, with a total abundance of 21%. Klitgaard (1995) reports a similar percent abundance of polychaetes (26%) within eleven demosponges. Polychaetes take advantage of the micro-niches found in sponge tissue, such as those found exclusively within the canals of the aquiferous system of the demosponge, *Anomoianthella lamella* (Magnino et al. 1999). Although collections were not done for every season in every reef, a similar fauna composition was found across an entire year of sampling among all reefs. Thus, *C. lutea* offers suitable habitat year-round for a mildly diverse group of facultative and ephemeral associated fauna.

Along with the infauna analysis, several other ecological and natural history observations were made. An unknown species of juvenile fish was observed taking shelter within the folds of *C. lutea* (Fig. 1b). Sponges providing shelter and refuge for fish have been reported before, but never for a calcareous sponge. Frith (1976) observed in several occasions the fish *Pholis gunnellus* and *Labrus bergylta*, laying within the folds of the sponge *Halichondria* (*Halichondria*) *panicea*. During collections, isopods were also observed scattering through the pores as the sponge was lightly disturbed prior to sampling.

A noteworthy observation, was that of a sea anemone inhabiting the surface of *C. lutea* (Fig. 1c). This represents the first record of an anemone, belonging to the order: Actiniaria, in association with a calcareous sponge (Fig. 2b). This anemone more than likely belongs to the genus *Bunodeopsis* (pers. comm. Estefania Rodriguez & Ricardo González-Muñoz), and has been previously found throughout the Caribbean growing on the leaves of the sea grasses, *Thalassia testudinum* and *Syringodium filiforme* (Day 1994; González-Muñoz et al. 2012), as well as in the demosponge *Aplysina cauliformis* (pers. comm. Deborah Gochfeld). Pearse (1950) also found anemones attached to the Caribbean demosponges *Ircinia strobilina* and *Amphimedon compressa*. All other published reports of sea anemones of the order Actiniaria have been in association with deep water hexactinellid sponges (Reiswig et al. 2011; Sanamyan et al. 2012; pers. comm. Christine Morrow, Christopher Kelley).

Despite its cryptic nature, the high number of associates within *C. lutea* emphasizes the key role and importance that marine sponges play in the ecosystem, especially since they are predicted to be the most dominant taxa on Caribbean reefs in the future (Pawlik et al. 2013; Loh et al. 2015). Within the Phylum Porifera, calcareous sponges tend to be, for various reasons, the most neglected group by sponge biologists (van Soest et al. 2012; van Soest and de Voogd 2015). We believe that further investigations of the class Calcarea will yield greater insights into reef biodiversity and the complex species interactions which unravel on the benthos (i.e., Ribeiro et al. 2016). These biotic interactions are especially important to observe and characterize now, since Caribbean and worldwide coral reefs are experiencing drastic community phase shifts due to a myriad of natural and anthropogenic stressors.

## ACKNOWLEDGMENTS

We would like to thank Michelle Klautau for her expert advice on calcareous sponges and for her constructive comments throughout the stages of this manuscript. Deborah Gochfeld, Ricardo González-Muñoz, Christopher Kelley, Christine Morrow, Henry Reiswig, Estefania Rodriguez, and Konstantin Tabachnick, for providing valuable information on sponge-anemone associations. Authors also thank Dr. Vasilis Gerovasileiou and an anonymous reviewer for their constructive comments, which greatly improved the manuscript. We would also like to thank the Department of Marine Science at the University of Puerto Rico - Mayagüez (UPRM-DMS) for continually providing resident researchers with small boat access for sampling across the La Parguera Natural Reserve area. Sponge collection permit was granted by The Department of Natural Resources Puerto Rico (Permit #O-VS-PVS15-MA-00021-22122015).

---

**Supplementary Table 1.** Ecological data for *Clathrina lutea* collected within this investigation, all reef sites are located within the Marine Reserve of La Parguera, southwest Puerto Rico.

| Sponge ID | Reef Site | Month | Depth (m) | Wet weight (g) |
| --- | --- | --- | --- | --- |
| CPR-4 | San Cristobal | April (2014) | 11.9 | 0.438 |
| CPR-5 | San Cristobal | April (2014) | 14.1 | 1.19 |
| CPR-6 | San Cristobal | April (2014) | 11.9 | 1.43 |
| CPR-12 | Turumote | May (2014) | 14.3 | 1.96 |
| CPR-15 | Turumote | May (2014) | 17.4 | 0.348 |
| CPR-10 | Enrique | May (2014) | 11.3 | 2.37 |
| CPR-19 | Turumote | June (2014) | 13.4 | 0.975 |
| CPR-200 | Enrique | July (2014) | 13.4 | 0.121 |
| CPR-25 | Turumote | August (2014) | 11.9 | 4.74 |
| CPR-29 | Media Luna | August (2014) | 17.9 | 4.53 |
| CPR-30 | Media Luna | August (2014) | 15.9 | 3.13 |
| CPR-33 | San Cristobal | August (2014) | 13.1 | 6.04 |
| CPR-36 | San Cristobal | August (2014) | 14.9 | 4.66 |
| CPR-37 | San Cristobal | August (2014) | 13.4 | 2.57 |
| CPR-42 | Turumote | September (2014) | 13.4 | 3.45 |
| CPR-47 | Turumote | September (2014) | 17.1 | 4.87 |
| CPR-58 | Mario | September (2014) | 14.3 | 3.49 |
| CPR-59 | Mario | September (2014) | 16.2 | 3.94 |
| CPR-61 | Mario | September (2014) | 17.1 | 3.06 |
| CPR-32 | San Cristobal | September (2014) | 12.5 | 2.39 |
| CPR-76 | Turumote | October (2014) | 13.1 | 2.31 |
| CPR-80 | Mario | October (2014) | 11.3 | 2.92 |
| CPR-69 | Margarita | October (2014) | 16.8 | 1.13 |
| CPR-72 | Margarita | October (2014) | 16.2 | 4.78 |
| CPR-87 | Pinnacles | November (2014) | 10.9 | 1.71 |
| CPR-88 | Pinnacles | November (2014) | 11.6 | 2.43 |
| CPR-94 | Pinnacles | November (2014) | 16.5 | 4.96 |
| CPR-108 | Pinnacles | December (2014) | 11.3 | 0.461 |
| CPR-110 | Pinnacles | December (2014) | 10.4 | 0.506 |
| CPR-113 | Pinnacles | December (2014) | 7.62 | 0.681 |
| CPR-123 | San Cristobal | January (2015) | 13.1 | 0.623 |
| CPR-124 | Pinnacles | January (2015) | 11.6 | 0.409 |
| CPR-130 | San Cristobal | February (2015) | 12.5 | 3.07 |
| CPR-126 | Pinnacles | February (2015) | 17.1 | 0.687 |
| CPR-127 | Pinnacles | February (2015) | 17.7 | 0.882 |
| CPR-137 | Turumote | March (2015) | 16.8 | 2.65 |
| CPR-139 | Turumote | March (2015) | 14.6 | 1.58 |
| CPR-140 | Turumote | March (2015) | 12.5 | 1.54 |
| CPR-142 | Laurel | March (2015) | 15.2 | 1.07 |
| CPR-154 | Turumote | April (2015) | 13.7 | 2.00 |
| CPR-155 | Turumote | April (2015) | 13.1 | 3.53 |
| CPR-156 | Media Luna | April (2015) | 14.9 | 3.75 |
| CPR-146 | Laurel | April (2015) | 11.9 | 5.28 |

**Supplementary Figure 1.**
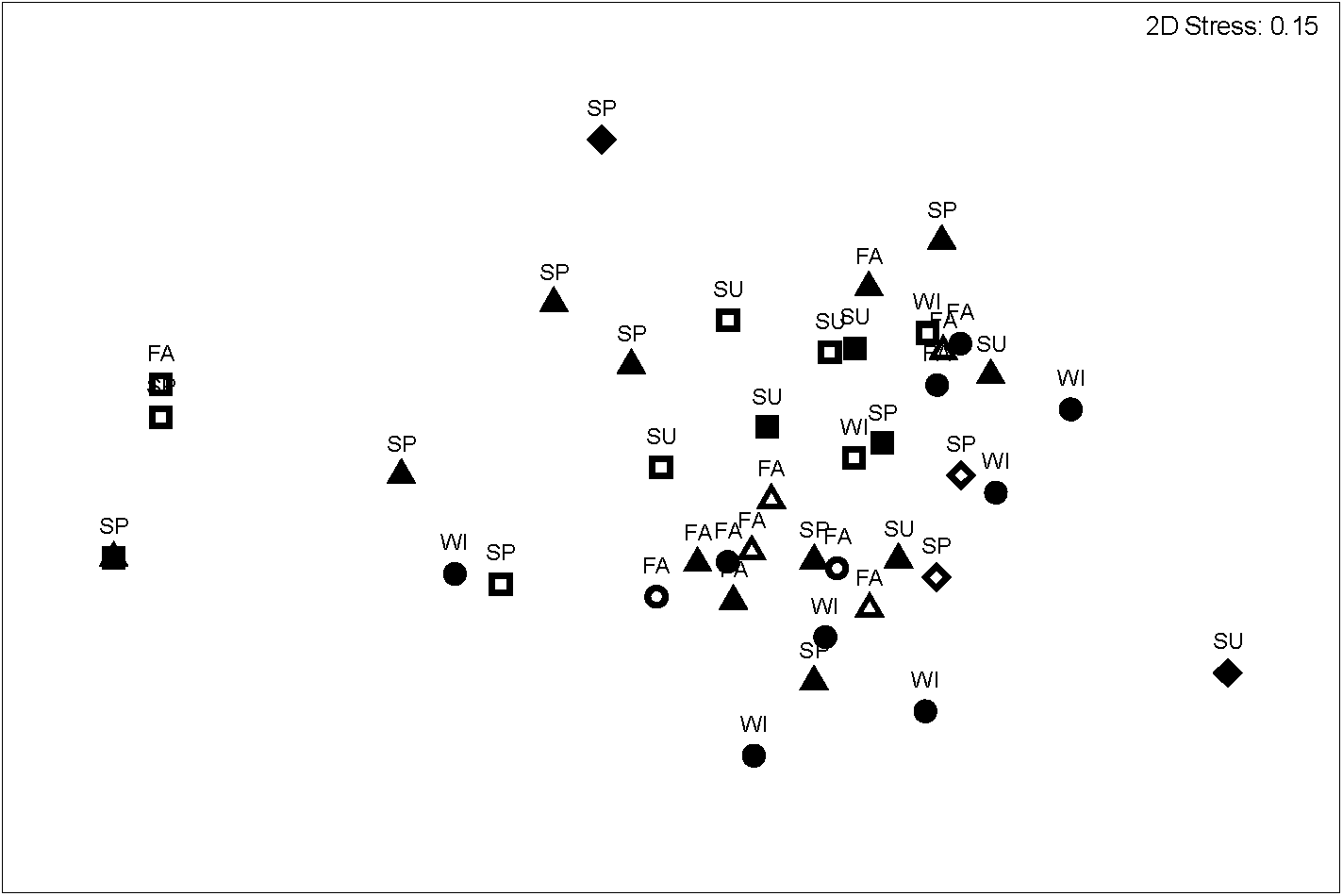
nMDS of Bray-Curtis similarities of square-rooted transformed data of assemblages associated with *Clathrina lutea* from eight different reefs at La Parguera Natural Reserve, PR (▴ = Turrumote, ▵ = Mario, ◼ = Media Luna, ◻ = San Cristobal, • = Pinnacles, **○** = Margaritas, ♦ = Enrique, ◊ = Laurel); sampled across different seasons (SP = Spring, SU = Summer, FA = Fall, WI = Winter).

